# Eosinophil peroxidase induces inflammation in a mouse model of dermatitis

**DOI:** 10.1101/2020.11.25.399006

**Authors:** Quinn R. Roth-Carter, Huijun Luo, Sergei I. Ochkur, Elizabeth A. Jacobsen, Meredith Datena, Allison D. Fryer, James J. Lee, David B. Jacoby

## Abstract

Eosinophils play an important role in mediating itch and inflammation in dermatitis. The role of the eosinophil granule protein eosinophil peroxidase (EPX) in mediating inflammation and itch was tested in a dermatitis mouse model. Mice were sensitized to trimellitic anhydride (TMA) and subsequently challenged chronically on the ear to establish dermatitis. Loss of EPX (in EPX (-/-) mice) or blocking EPX with the drug resorcinol significantly reduced dermatitis in mice exposed to TMA. Resorcinol also reduced levels of thymic stromal lymphopoietin protein (TSLP) in skin. Further studies showed that EPX increased different cytokines in keratinocytes in cell culture via two distinct mechanisms. EPX induced TSLP expression requires lysophosphatidic acid signaling while EPX induced expression of TNF-α, CSF2, CSF3, and IL1α required IL-1 signaling. We also showed that blocking IL-1 reduced inflammation in skin following TMA exposure in mice. Thus, EPX is an important mediator of inflammation and itch, that are mediated via at least two pathways. This suggests that both EPX and its’ signaling pathways may provide novel therapeutic strategies in dermatitis.

## INTRODUCTION

Eosinophils were first observed in skin of patients with atopic dermatitis in 1975(Steigleder and Inderwisch 1975) and were subsequently also shown to be increased increased in peripheral blood (Uehara et al. 1990; Jenerowicz et al. 2007). Eosinophil granule proteins are increased in serum with atopic dermatitis, and their concentration correlates with severity of disease (Czech et al. 1992; Kagi et al. 1992; Halmerbauer et al. 1997; Taniuchi et al. 2001). However, despite strong associations between eosinophil presence and atopic dermatitis symptoms, their role in this skin disease is still unclear.

Eosinophils are important mediators of dermatitis in a mouse model that closely mimics the inflammatory response seen in the skin in patients with atopic dermatitis (Lee et al. 2015). Immunostaining of skin from patients with atopic dermatitis reveals release of the eosinophil peroxidase (EPX) (Foster et al. 2011), an eosinophil granule protein which has been implicated in causing inflammation in other disease settings (Forbes et al. 2004).

The epithelial cell derived cytokine thymic stromal lymphopoietin (TSLP) is elevated in skin in patients with atopic dermatitis, and over expression of TSLP in skin of mice leads to itch and dermatitis (Soumelis et al. 2002; Yoo et al. 2005; Sano et al. 2013). In addition to participating in inflammation (Soumelis et al. 2002; Watanabe et al. 2004; Gao et al. 2010; Sano et al. 2013; Luo et al. 2014) TSLP can also directly activate sensory nerves to cause itch (Wilson et al. 2013).

We have previously showed that eosinophils are important in mediating itch and inflammation in mice with dermatitis caused by sensitization and cutaneous application of trimellitic anhydride (TMA) (Lee et al. 2015). Here we show that eosinophil peroxidase (EPX) is central to these responses, and that EPX activity is required for increased TSLP expression in this model. We also show that EPX activates two distinct pathways to directly increase cytokine expression in keratinocytes. In one pathway, EPX increases TSLP expression by signaling through lysophosphatidic acid (LPA). In a second pathway, EPX increases expression of CSF2, CSF3, TNF and IL1α, by signaling through IL-1 receptors. Finally, we show that blocking IL-1 in a mouse model of dermatitis reduces inflammation, similar to blocking EPX activity.

## RESULTS

### Eosinophil peroxidase is an important mediator of itch and inflammation

To test the role of EPX in dermatitis we chronically exposed BABL/C and EPX -/-mice to TMA (add concentration and time-I know methods are later-but give your reader a clue) on one ear to induce inflammation and itching. The contralateral ear was used as a vehicle control. Chronic exposure of wildtype BALB/c mice to TMA significantly (?) increased scratching of the TMA treated ear and increased ear thickness (a general measure of inflammation), compared to the contralateral control ear (Figure 1A-B). EPX^-/-^ mice had significantly reduced scratching compared to WT mice (Figure 1A), and a significant reduction in ear thickness in the TMA treated ear compared to WT mice (Figure 1B). TMA also caused a robust recruitment of eosinophils into the skin in WT mice, while EPX^-/-^ mice had reduced eosinophils in skin in TMA treated ears compared to TMA treated ears from WT mice (Figure 1E). This effect was selective for EPX, as mice that lack another eosinophil granule protein, major basic protein, had no change in itch or increase in ear thickness in response to TMA compared to WT mice (see Figure S1).

**Figure 1.**
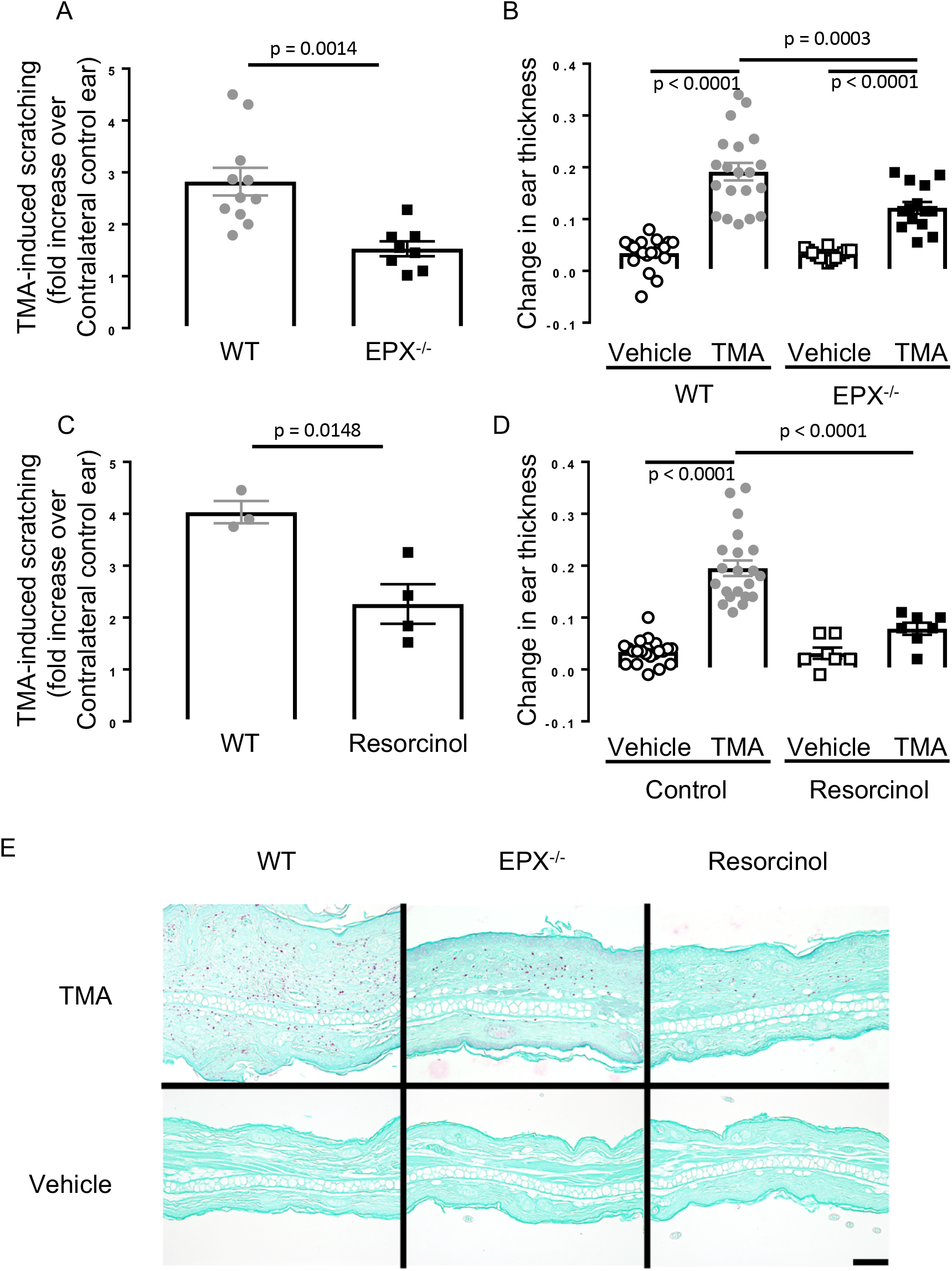
EPX is required for inflammation and itch caused by exposure to TMA

To further test the role of EPX in mediating dermatitis, mice were treated with a peroxidase inhibitor, resorcinol (1.25 mg/kg) (Forbes et al. 2004; Chu et al. 2014). Resorcinol dose dependently inhibited the activity of EPX in a peroxidase activity assay (Figure S2). Mice treated with resorcinol had significantly reduced ear scratching compared to vehicle treated mice (Figure 1C). Resorcinol also significantly inhibited TMA-induced increased ear thickness (Figure 1D). Finally, resorcinol reduced eosinophils present in skin in TMA treated ears (Figure 1E). Reducing TMA induced ear scratching and increased ear thickness, as well as decreasing skin eosinophils, suggests that EPX is an important mediator of TMA induced itch and inflammation in skin, and that this is mediated by its peroxidase activity.

### Resorcinol reduces inflammation and cytokine expression

EPX activity in TMA induced skin inflammation was tested in another strain of mice, C57BL/6 that is known to be more or less sensitive to (antigens/irritants/add how these are different from black6). As in BALB/c mice, chronic treatment of C57BL/6 mice with TMA increased ear thickness (Figure 2A) increased eosinophil numbers in skin (Figure 2B), and increased TSLP in ears (Figure 2C). Resorcinol significantly reduced all three parameters, confirming that TMA induced ear itch and local inflammation are likely mediated by EPX and TSLP (Figure 2A-C) in C57BL/6 similarly to BALB/c mice.

**Figure 2.**
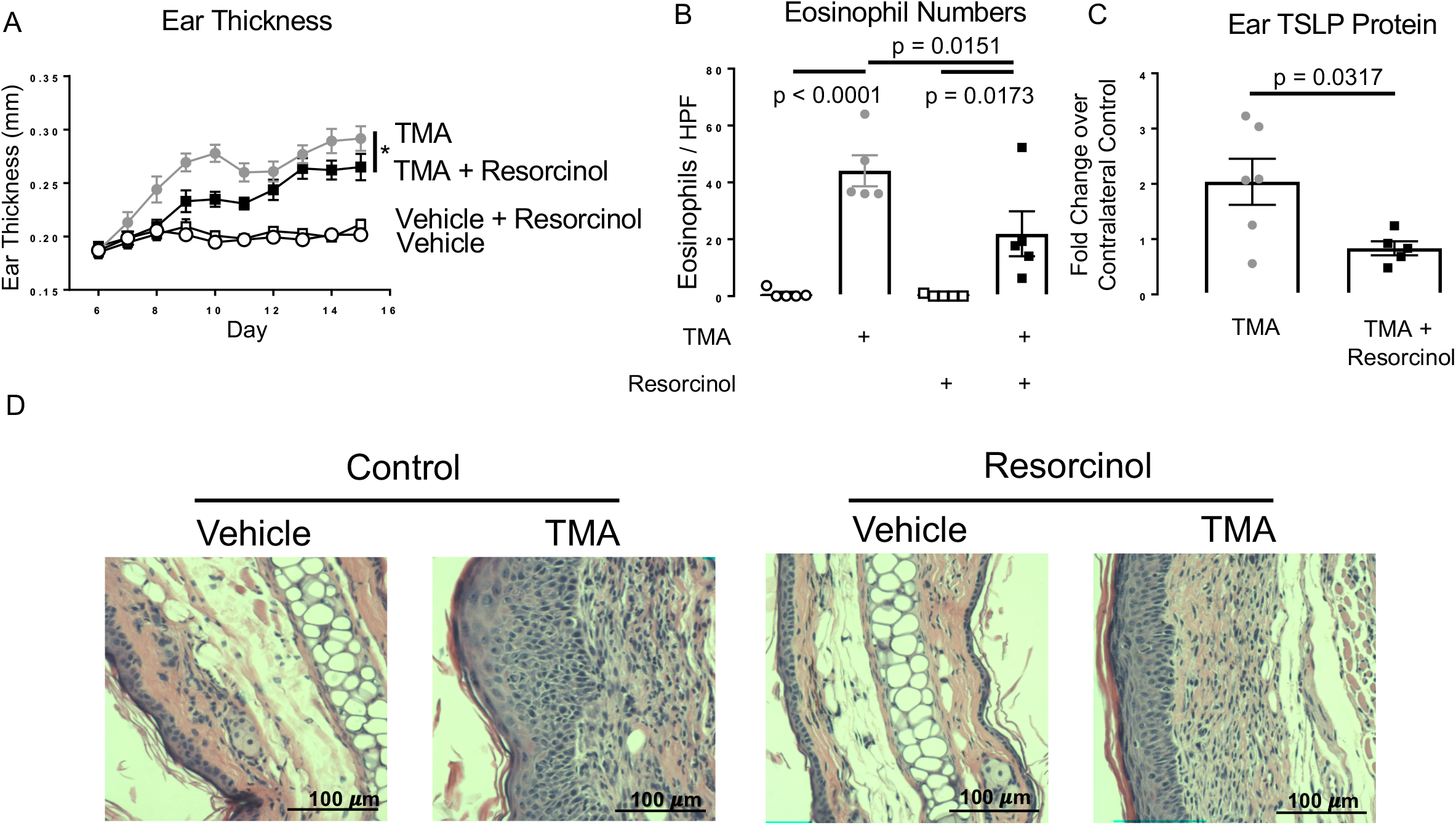
EPX is required for inflammation and the increase in TSLP caused by exposure to TMA

### Eosinophil peroxidase increases cytokine expression in keratinocytes

The effect of EPX activity on cytokine gene expression in keratinocytes was measured using a common cytokine profiler array. Mouse keratinocytes were treated with purified human EPX, with or without its substrates H_2_O_2_ and Br^-^. EPX plus substrates increased expression of TSLP, TNF, CSF2, CSF3, and IL1α (Figure 3A, Table S1). None of the substrates alone, or EPX without substrates, increased gene expression for these cytokines (Figure 3A). EPX activity was required for increased cytokine expression as both heat inactivation (80°C for 5 min) and the EPX inhibitor resorcinol (30 μM) prevented expression (Figure 3B-C). EPX plus substrates did not affect cell viability as measured by MTT assay (data not shown).

**Figure 3.**
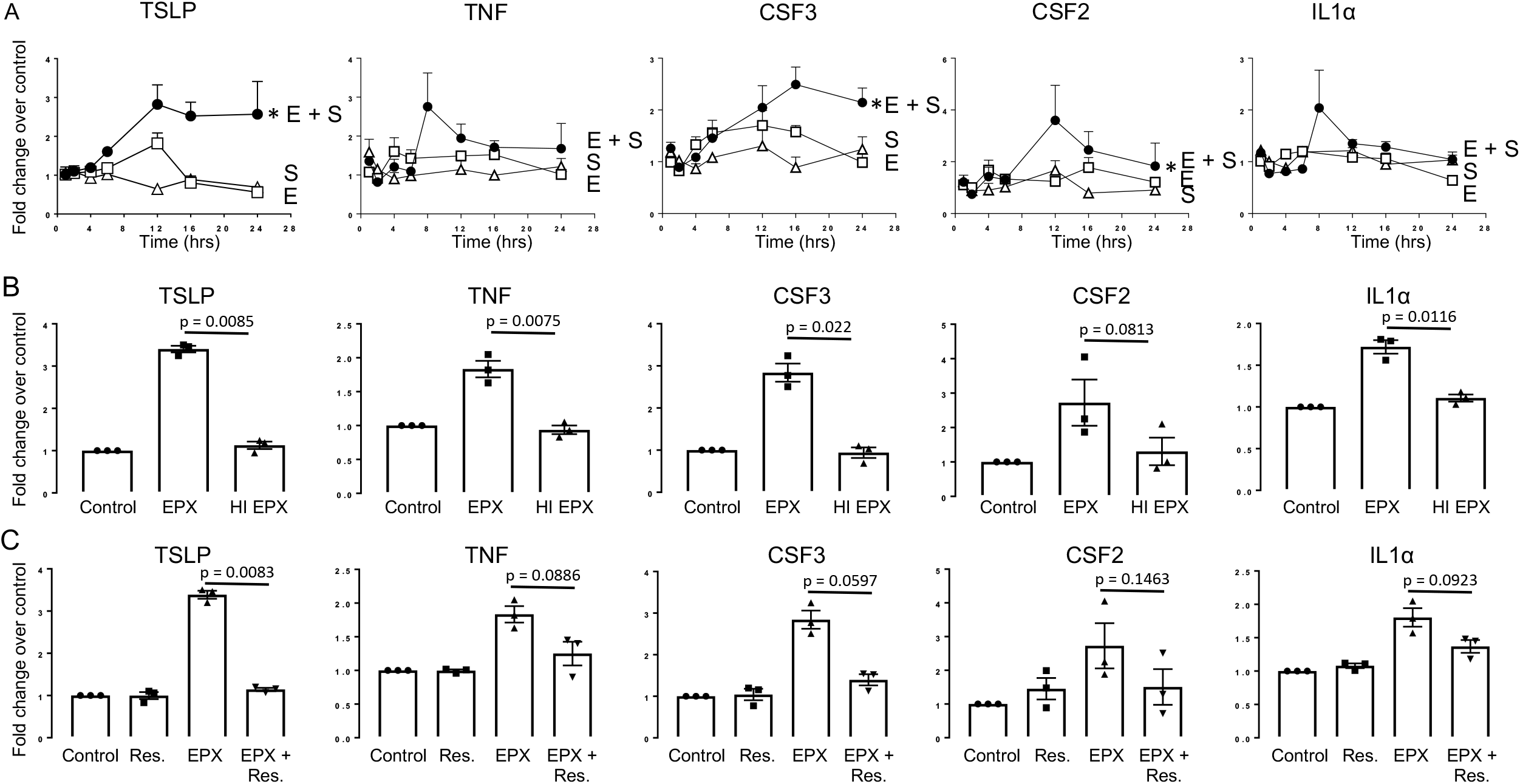
EPX increase cytokine expression in keratinocytes

### EPX increases cytokine expression in keratinocytes through a soluble mediator

While EPX increased gene expression of cytokines in isolated keratinocytes, this increase in expression was delayed 8 - 12 hours after treatment (Figure 3A). To explain this delay, we hypothesized that this cytokine gene expression was mediated by an intermediate mediator and tested this using conditioned media taken from keratinocytes treated with EPX and its substrates. Conditioned media increased cytokine gene expression after only 4 hours (Figure S3).

### Lysophosphatidic acid (LPA) increases TSLP expression, but does not affect gene expression for other cytokines

LPA, which is known to increase TSLP expression in airway epithelial cells (Medoff et al. 2009), was tested as a potential mediator of increased cytokine expression in keratinocytes induced by EPX. mRNA for all six LPA receptors (LPAR 1-6) was detected in cultured keratinocytes (Figure 4A). We also tested keratinocytes for expression of secreted phospholipase A2 enzymes (sPLA2) enzymes, which are involved in production of LPA. Of 11 sPLA2 enzymes tested, sPLA2G2F, sPLA2G7, sPLA2G2E, and sPLA2G12A were expressed (Figure 4B). Thus, keratinocytes express both, receptors to respond to LPA, and enzymes that produce LPA. Treating keratinocytes with LPA dose dependently and significantly increased TSLP gene expression in keratinocytes (Figure 4C). This effect was selective for TSLP, as LPA did not affect expression of any other measured cytokine (Figure 4D).

**Figure 4.**
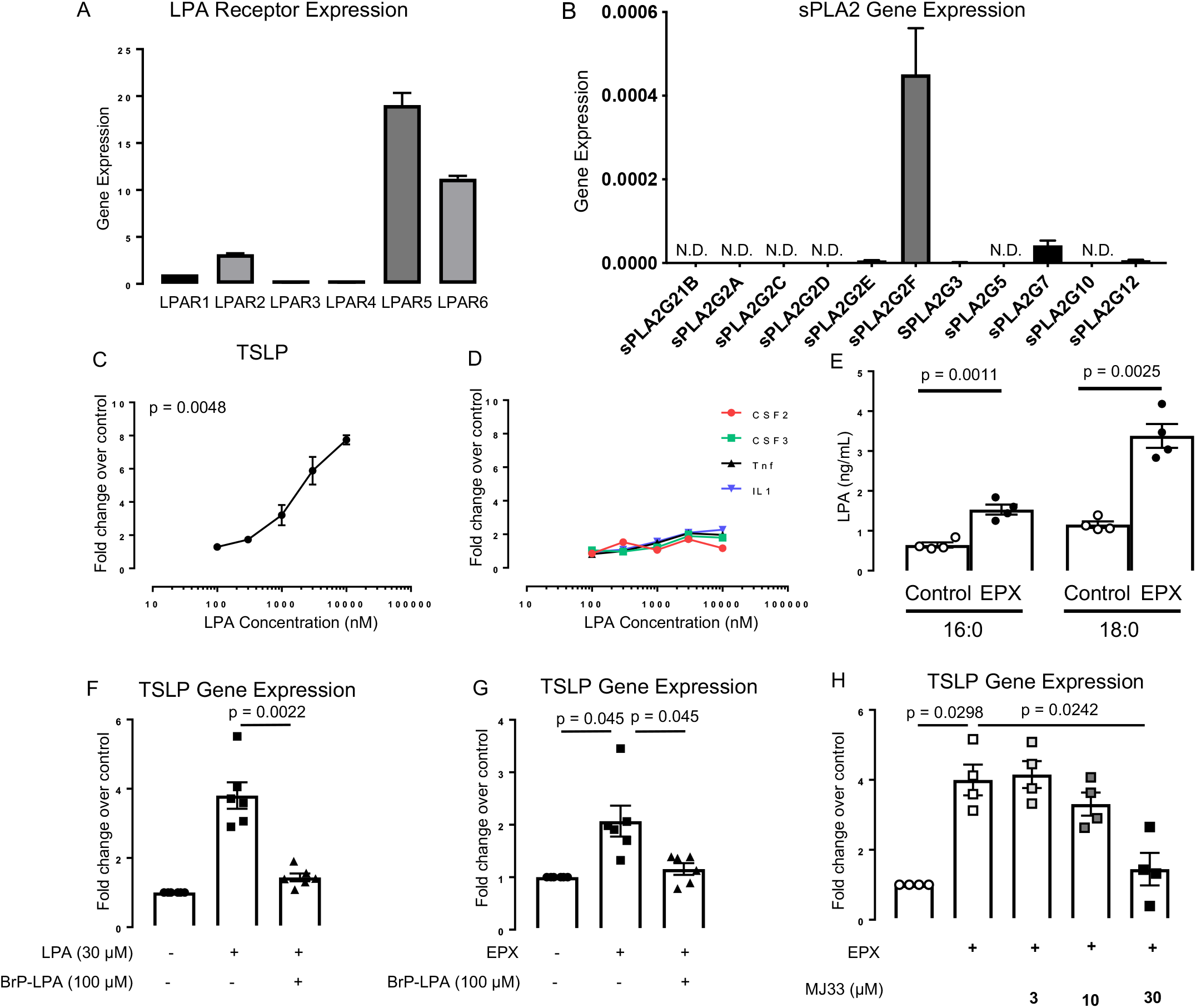
LPA mediates the increase in TSLP expression caused by EPX

### Lysophosphatidic acid production is required to for EPX to stimulate TSLP expression in keratinocytes

Two species of LPA (16:0 and 18:0 LPA) were increased in cell culture media of mouse keratinocytes treated with EPX and substrates compared to media from untreated keratinocytes (Figure 4E). Treating keratinocytes with the pan-LPA receptor antagonist BrP-LPA (100μM) inhibited the increase in TSLP expression caused by LPA (Figure 4F), and prevented the increase in TSLP gene expression in keratinocytes treated with EPX and its substrates (Figure 4G). Preventing generation of LPA using the sPLA2 inhibitor MJ33 also dose dependently inhibited the increased TSLP expression in keratinocytes exposed to EPX and its substrates (Figure 4H). MJ33 did not affect keratinocyte viability (data not shown).

### IL-1 signaling is required for EPX to increase cytokine expression, except for TSLP

While LPA signaling plays a key role in EPX induced increased TSLP expression in keratinocytes, LPA did not stimulate expression of other cytokines that were increased by EPX (Figure 4D). Conversely, treating keratinocytes with IL-1α significantly increased gene expression for CSF2, CSF3, TNF and IL-1α in keratinocytes, but did not increase TSLP gene expression (Figure 5A-E). Treating keratinocytes with the IL-1 receptor antagonist anakinra (30 μg/mL) prevented the EPX induced increase in gene expression of CSF2, CSF3, TNF, and IL-1α in keratinocytes (Figure 5H-J), but did not prevent the EPX induced increase TSLP expression (Figure 5F).

**Figure 5.**
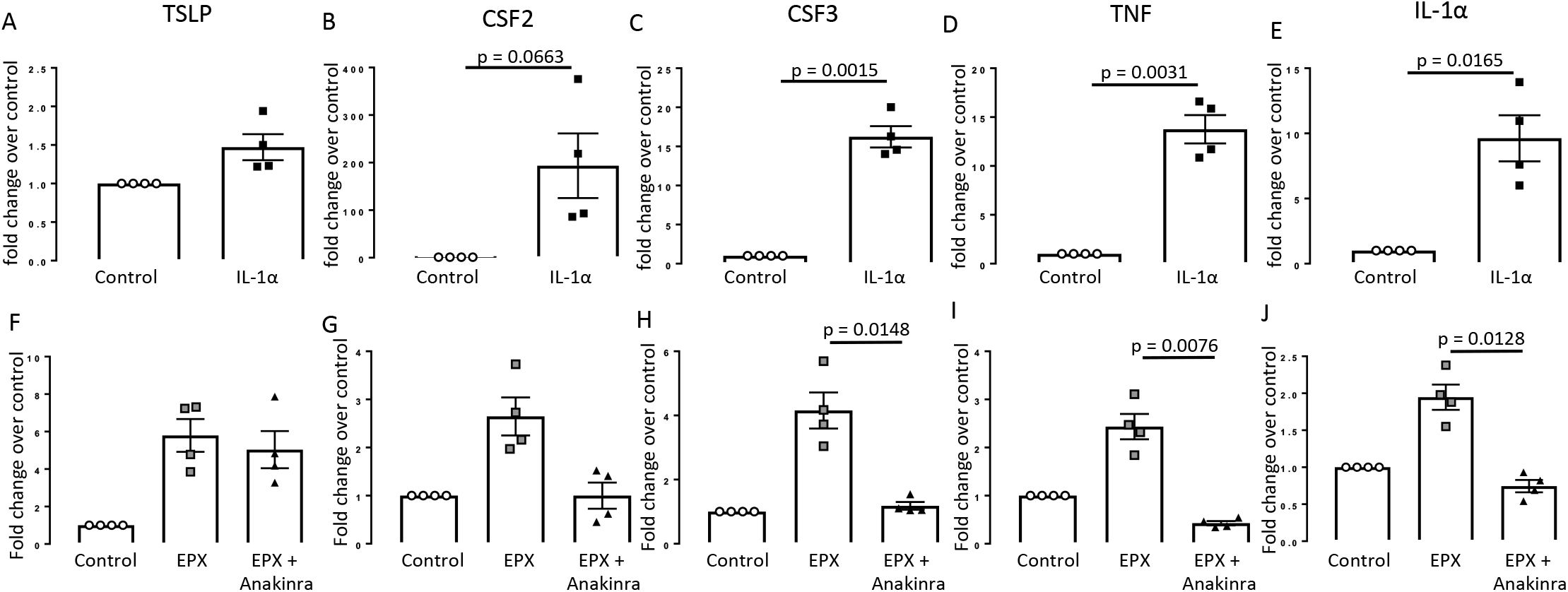
IL-1 mediates the increase in cytokine expression caused by EPX

### Anakinra reduces skin inflammation in vivo

To determine whether IL-1 was critical for inflammation in our animal dermatitis model, C57BL/6 mice were treated with the IL-1 receptor antagonist, anakinra (30mg/kg, i.p.) daily during TMA challenge. Anakinra significantly inhibited the increase in ear thickness (Figure 6A), the number of eosinophils present in skin (Figure 6B), and the epidermal thickening caused by TMA (Figure 6C).

**Figure 6.**
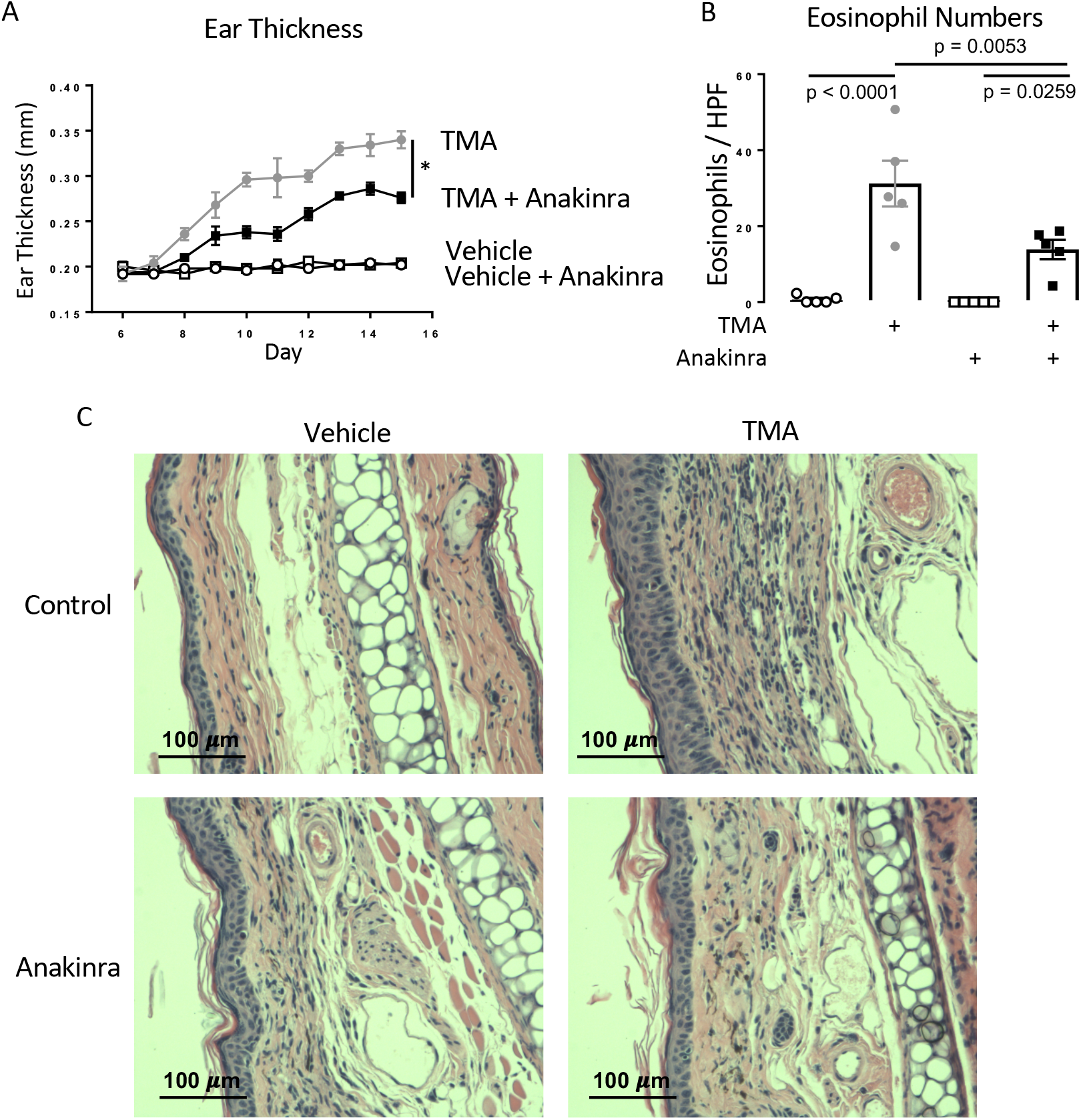
IL-1 is an important mediator of inflammation in a chronic contact dermatitis model.

## DISCUSSION

Here, we show that EPX is an important mediator of inflammation and itch in an animal model of dermatitis in two strains of mice. EPX activity also increases cutaneous TSLP in mouse skin. EPX, with its substrates, increased TSLP, CSF3, CSF2, TNF and IL-1α gene expression in keratinocytes. This effect required peroxidase activity of EPX, as the peroxidase inhibitor resorcinol blocked the increase in cytokine gene expression caused by EPX, as did heat inactivation of EPX. This closely matches in vivo data where inhibiting EPX reduced skin inflammation and TSLP levels.

EPX increases expression of these cytokines in keratinocytes using two pathways. For TSLP, EPX drives production of LPA which then increases TSLP expression in keratinocytes, and this could be blocked by either preventing LPA production with the sPLA2 inhibitor MJ33 or by blocking LPA receptors with the antagonist BrP-LPA. In contrast, for CSF3, CSF2, TNF and IL-1α, EPX increased gene expression through IL-1, which could be blocked with the IL-1 receptor antagonist anakinra. Moreover, in our mouse model, blocking IL-1 with anakinra or blocking EPX with resorcinol each reduced inflammation to a similar level. These data support other recent work that identified IL-1 as an important mediator of inflammation in atopic dermatitis (Kezic et al. 2012; Bernard et al. 2017). This work also identifies LPA as a potential new mediator and therapeutic target in atopic dermatitis.

Although LPA has previously been shown to increase TSLP expression in airway epithelial cells (Medoff et al. 2009), this is the first evidence that LPA increases TSLP expression in keratinocytes. Whether LPA levels are also increased in skin of patients with atopic dermatitis is unknown. However, the levels of autotaxin, an enzyme involved in LPA production, are elevated in serum of patients with atopic dermatitis, and correlate with disease severity (Nakao et al. 2014). Like TSLP, LPA has also been shown to directly activate sensory nerves and stimulate itch (Kittaka et al. 2017), suggesting redundancy in mechanisms causing itch.

Our group showed that eosinophils are an important mediator of itch in a mouse model of dermatitis (Lee et al. 2015), and we have further shown here that EPX is important for itch. Based on the observation that both TSLP and LPA can activate sensory nerves to cause itch (Wilson et al. 2013; Kittaka et al. 2017), we propose that this is a pathway by which eosinophils, and specifically EPX, can cause itch in this model. A number of previous studies suggest a role for this mechanism in atopic dermatitis. TSLP levels are elevated in skin in patients with atopic dermatitis (Soumelis et al. 2002; Sano et al. 2013). Staining for EPX in skin of patients with atopic dermatitis also reveals release of EPX (Foster et al. 2011). The link between EPX activity and TSLP in these experiments suggests that eosinophils increase TSLP expression in skin in patients with atopic dermatitis, potentially by stimulating production of LPA.

IL-1 is a well characterized mediator of inflammation known to be elevated in plasma of patients with atopic dermatitis (Thijs et al. 2017). Here we show that IL-1 is critical for EPX to increase gene expression of several cytokines (CSF3, CSF2, TNF and IL-1α) in keratinocytes, and that blocking IL-1 in vivo significantly reduces inflammation. All of these cytokines, CSF3, CSF2, TNF and Il-1α, have been found to be elevated in plasma of patients with atopic dermatitis (Thijs et al. 2017). CSF3 (GM-CSF) is a growth factor for inflammatory cells, including eosinophils (Burstein et al. 1992) and also primes and activates eosinophils (Tai and Spry 1990). TNF was recently shown by our group to stimulate eosinophilopoiesis after exposure to ozone (Wicher et al. 2017), and increases expression of eotaxin in epithelial cells, a potent chemokine for eosinophils (Lilly et al. 1997). IL-1 also directly stimulates expression of adhesion molecules on endothelial cells that eosinophils use to migrate into tissue (Eissner et al. 1997).

While we propose that LPA and TSLP are important inducers of itch, TSLP may affect inflammation as well. TSLP stimulates eosinophil degranulation in vitro (Cook et al. 2012), increases eosinophilopoiesis (Denburg et al. 2015), and increases eosinophil survival (Wong et al. 2010). We hypothesize that EPX activity initiates a feedforward loop, though Il-1 and LPA, that recruits and activates more eosinophils, leading to chronic inflammation.

The mechanism by which EPX peroxidase activity stimulates production of LPA and IL-1 has not been determined. EPX uses hydrogen peroxide to oxidize bromide to produce hypobromous acid (van Dalen and Kettle 2001), which is a potent oxidizing agent that is toxic to both parasites and human cells(Klebanoff et al. 1989; Slungaard and Mahoney 1991). Our findings show that EPX can activate signaling pathways without triggering cell death, as these treatments did not have any effect on cell viability. While products of EPX are toxic at high concentrations, at low concentrations these products may serve as signaling molecules.

These data demonstrate that EPX is in an important mediator of itch and inflammation in an animal model of dermatitis. Similar interactions between eosinophils and epithelial cells may also be present in the airways in asthma and in the esophagus in eosinophilic esophagitis, diseases that also have increased levels of eosinophils and TSLP in tissue. Data presented here describe a mechanism by which eosinophils, through eosinophil peroxidase, can promote chronic inflammation in tissue, and identify novel pathways that can be targeted to reduce symptoms in atopic dermatitis and be tested in other chronic inflammatory diseases.

## METHODS

### Mice

All studies were performed with 8-14 week-old mice on a BALB/c or C57BL/6 background. EPX^-/-^ mice (Denzler et al. 2001) were bred at the Mayo Clinic Arizona animal facility on a BALB/c background. BALB/c control mice were either bred in house or purchased directly from Jackson Laboratories (Jackson Research Laboratories, Bar Harbor, Maine). C57BL/6 wild type mice were bred in the OHSU animal facility. All protocols and studies involving animals were performed in accordance with guidelines of National Institute of Health and approved both by the Institutional Animal Care and Use Committee of Mayo Foundation for Medical Education and Research and by Institutional Animal Care and Use Committee of Oregon Health & Sciences University. Each experimental group included 5-14 mice, and as dermatitis was induced on only one ear, the contralateral unaffected ear served as a control.

### Mouse dermatitis model

Mice were sensitized and challenged to produce dermatitis as previously described(Schneider et al. 2009; Lee et al. 2015). Briefly, 50 μL of 5% TMA (Sigma) solution in 4:1 acetone/olive oil was applied to a shaved portion of the back on days 0 and 5 to sensitize the animals. Mice were challenged daily by application of 15 μL 2% TMA in acetone/olive oil vehicle to the right ear on days 6 – 14. Contralateral control ears were treated with vehicle. Mice were anesthetized with isoflurane before every sensitization and challenge. Thickness of both ears was measured before each challenge on days 6-14 or once on day 14? using a digital caliper (Mitutoyo). Animals were single housed throughout the sensitization and challenge protocol to prevent cross contamination of TMA solution. Mice were treated with either resorcinol (1.25 mg/kg, Sigma), anakinra (30mg/kg, Amgen), or vehicle (PBS) before each challenge. On day 15, mice were euthanized by i.p. pentabarbitol (300 mg/kg), and ears were removed, flash frozen in liquid nitrogen and stored at −80^°^C until analyzed by ELISA. For histology samples, ears were removed and fixed for 16 hours in Zamboni’s fixative.

### Assessment and quantification of itching in response to TMA exposure

On day 14, the last day of challenge, a high definition time-lapse video system was used to record scratching (Lee et al. 2015). Scratching events were counted manually by 3 independent investigators who were blinded to mouse genotype and drug treatment. At least 100 total scratching events, defined as deliberate scratching of the ear in question using a hind paw, were counted from each video. Itching scores associated with TMA exposure for each mouse within a given genotype or treatment were assessed as fold-increase of ear scratching events over the contralateral control ear of that mouse.

### TSLP measurements

Ears were homogenized (Tissue Terror, Biospec Products) in protein extraction buffer (10 mM Tris-HCL, pH 7.5; 0.5 mM EDTA; 0.5 mM EGTA; 1% Triton X-100; 0.5 mM PMSF; and Protease Inhibitor Cocktail [Thermo Fisher Scientific]). Protein concentration was measured using a BCA protein assay (Thermo Fisher Scientific). TSLP levels were measured by ELISA (R & D Systems), normalized to total protein, and expressed as a fold change of TMA treated ear over the contralateral control ear.

### Eosinophil quantification

Mouse ears were paraffin embedded, sectioned and stained with hematoxylin and eosin. Eosinophils were counted by a blinded observer. 3 random high-powered fields were selected from each slide, and total eosinophils were counted in each field. Average number of eosinophils across the fields were calculated for each sample.

### Immunohistochemistry

Rat anti-mouse eosinophil associated ribonuclease monoclonal antibody (MT-32.1.3, 1:500) was used to label eosinophils in mouse skin for immunohistochemistry (Doyle et al. 2013).

### Cell culture

Keratinocytes were isolated from mouse tail skin using modifications of previous protocols (Lichti et al. 2008). C57BL/6 mice were euthanized by pentobarbital (300 mg/kg, i.p.). Tails were removed and skin was pulled from the tail. The skin was floated dermis side down on 0.25% trypsin (Gibco) overnight at 4^°^ C, and the following day the dermis was peeled from the epidermis. The epidermis was minced and triturated in Minimum Essential Media (MEM, Gibco) with 5% Fetal Bovine Serum (FBS, Hyclone), penicillin, streptomycin and amphotericin B (PSF, Thermo Fisher Scientific). Disaggregated keratinocytes were passed through a 100 μm cell strainer (Corning). Keratinocytes were plated onto fibronectin (from human plasma, Sigma) and collagen (bovine type 1, Sigma) coated plates in Keratinocyte Growth Medium 2 (KGM-2, Lonza) at a density of 2 x 10^5^ cells/cm^2^ and cultured at 37^°^C at 5% CO_2_. Once cells reached confluence they were cultured in KGM-2 lacking hydrocortisone for 24 hours.

Keratinocytes were treated with purified human EPX (30 nM, Provided by Dr. Gerald Gleich, University of Utah(Agosti et al. 1987)), with or without its substrates hydrogen peroxide (100 μM, Fisher Chemical) and sodium bromide (100 μM). Drugs and doses used: resorcinol (30 μM, Sigma), anakinra (30 μg/mL, Amgen), MJ33 (Sigma, 3 – 30 □M), BrP-LPA (100 μM, Echelon), lysophosphatidic acid (Polar Biolipids), and rmIL-1 *α* (3 ng/mL, Peprotech).

### mRNA isolation and quantitative reverse transcriptase PCR

RNA was isolated from keratinocytes cultures using a RNeasy Kit (Qiagen). cDNA was prepared using 100 ng of total RNA by reverse transcription using Superscript III (Invitrogen). Gene expression was measured using primers in Table 1. qRT-PCR was performed using SYBR Green (Qiagen) assay in Applied Biosystems 5500 Fast Thermocycler. Changes in gene expression were normalized to 18S rRNA, and relative quantities of RNA were determined using the delta-delta CT method (Schmittgen and Livak 2008).

### Measurement of LPA

LPA levels in cell culture supernatants were measured by lipid chromatography/mass spectroscopy using Applied BioSystems 4000 QTRAP. Lipids were extracted using methanol, and samples were then dried and reconstituted before analysis. LPA concentrations were then determined by comparison to a standard curve using purified lipids (Avanti Polar Lipids).

### Data Analysis

All data are expressed as mean ± SEM. Comparisons between two groups were done using t-test. Multiple comparisons were done using one-way ANOVA with Tukey post-hoc test to determine differences between groups. For experiments comparing ear thickness over the course of experiment, groups were compared using two-way ANOVA for repeated measures. A p value of less 0.05 was considered statistically significant.

## Acknowledgments

The authors would like to thank Dr. Gerald Gleich for providing purified eosinophil peroxidase for these experiments. We would also like to thank Dr. Becky Proskocil for aid in editing the manuscript and editing figures. This work was funded by grants from the National Institute of Arthritis and Musculoskeletal and Skin Diseases, National Institute of Heart, Lung and Blood, and the National Cancer Institute. National Institute of Heart, Lung and Blood HL124165 and National Institute of Arthritis and Musculoskeletal and Skin Diseases AR061567 was awarded to DBJ and JJL. QRRC was supported by T32HL083808 and T32CA106195.

## Author Contributions

Conceptualization: QRC, HL, EAJ, ADF, JJL, DBJ; Formal Analysis: QRC, HL; Funding Acquisition: ADF, JJL, DBJ; Investigation; QRC, HL, SIO, MD; Visualization: QRC, HL: Writing - Original Draft Preparation: QRC; Writing - Reviewing and Editing: HL, SIO, EAJ, MD, ADF, DBJ

---

**Figure 1. Eosinophil peroxidase is a mediator of itch and inflammation in a mouse model of dermatitis**. A) TMA induced itching was measured using time lapsed videography after the final TMA exposure. Quantification of scratching of TMA treated ears in WT and EPX^-/-^ mice were compared to the vehicle treated contralateral control ear from the same mouse. Data is expressed as a fold increase in scratching of TMA treated ear over contralateral control ear from same mouse. B) Sensitized WT and EPX ^-/-^ mice were challenged on the ear with TMA, and ear thickness was measured on day 6 before the first challenge and on day 15. Change in ear thickness is expressed as difference in mm between two readings. C) TMA induced ear scratching quantified as described above. Mice were treated with either resorcinol daily (1.25 mg/kg i.p.) or vehicle just before challenge on the ears with TMA. D) Ear thickness was measured as described above in mice treated with either resorcinol daily just before challenge or vehicle as control. E) Eosinophils were labeled in skin by staining eosinophils pink with a rat anti-eosinophil associated ribonuclease antibody.

**Figure 2. Resorcinol reduces inflammation and TSLP in a mouse model of dermatitis**. A) Sensitized mice were challenged with TMA on the ear, and contralateral control ears were challenged with vehicle as control. Mice were treated daily during challenge with either resorcinol (1.25 mg/kg i.p.) or vehicle as control. Ear thickness was measured just before each challenge using an electric caliper. Curves were compared using a two-way ANOVA with repeated measures, *p<0.05. B) Eosinophils were counted from hematoxylin and eosin stained samples. Data are expressed as average number of eosinophils per high powered field. C) TSLP levels were measured from protein extracted from ears and normalized to protein concentrations. The data are then expressed as a fold change of μg TSLP/mg protein in the TMA exposed ear over the contralateral control ear. D) Hematoxylin and eosin stained ear tissue sections from TMA exposed mice, and control exposed contralateral control ears.

**Figure 3. Eosinophil peroxidase increases cytokine expression in keratinocytes**. A) Cytokine gene expression in keratinocytes treated with EPX (30 nM), H_2_O_2_ (100 μM) and Br^-^ (100 μM) compared to control. Gene expression was measured at 1, 2, 4, 6, 8, 12, 16 and 24 hours after treatment. B) Cytokine gene expression in keratinocytes treated with EPX, plus substrates H_2_O_2_ and Br^-^ or heat inactivated (HI) EPX plus substrates. C) Cytokine gene expression in keratinocytes treated with resorcinol (30 nM), eosinophil peroxidase and its substrates. Data in all figures expressed as fold change in gene expression over control cells.

**Figure 4. Lysophosphatidic acid increases TSLP gene expression in keratinocytes**. A) Gene expression for all described LPA receptors was measured using real-time PCR in primary keratinocyte cell cultures and levels were normalized to 18s rRNA. B) Gene expression of for all described secreted phospholipase A2 enzymes was measured in primary keratinocytes using real-time PCR and normalized to 18s rRNA. ND= not detected. C-D) Cytokine gene expression in keratinocytes treated with increasing concentrations of lysophosphatidic acid for 4 hours. Data are expressed as fold change over control cells. E) 16:0 and 18:0 LPA levels in media of keratinocytes treated with EPX plus substrates for 8 hours were measured using LC-MS. Data are expressed as concentration of LPA in ng/mL. F) TSLP gene expression in keratinocytes treated with EPX plus substrates and the pan-LPA receptor antagonist BrP-LPA at 100 μM for 8 hours. Data are expressed as fold change over control cells. G) TSLP gene expression of keratinocytes treated with EPX plus substrates and the pan sPLA2 inhibitor MJ33 at noted concentrations for 8 hours. Data expressed as fold change over control cells.

**Figure 5. IL-1 is required for the increase in CSF2, CSF3, TNF, and IL-1a gene expression in keratinocytes**. A-E) Cytokine gene expression in keratinocytes treated with IL-1α for 2 hours. F-J) Cytokine gene expression in keratinocytes treated with EPX plus substrates with the IL-1 receptor antagonist Anakinra (30ug/mL) for 8 hours. Data for all figures expressed as fold change over control treated cells.

**Figure 6. Anakinra reduces inflammation in a mouse model of dermatitis**. A) Sensitized mice were challenged with TMA on the ear, and contralateral control ears were challenged with vehicle as control. Mice were treated daily during challenge with either anakinra (30 mg/kg i.p.) or vehicle as control. Ear thickness was measured just before each challenge using an electric caliper. B) Eosinophils were counted from hematoxylin and eosin stained samples. Data are expressed as average number of eosinophils per high powered field. C) Hematoxylin and Eosin stained ear tissue sections from TMA exposed mice, and control exposed contralateral control ears. *p<0.05.

