## Supplemental Material for "Eosinophil peroxidase induces inflammation in a mouse model of dermatitis"

**Online Repository**

**Methods**

**Mice**

Major basic protein knockout ($MBP^{-/-}$) mice were bred at the Mayo Clinic Arizona Facility on a Balb/C background[1]. Each experimental group contains 4-14 mice.

**Qiagen RT2 profiler array**

RNA was isolated from keratinocytes cultures using a RNeasy Kit (Qiagen). cDNA was prepared by reverse transcription using Superscript III (Invitrogen). Gene expression was measured using the common cytokines RT2 profiler array (Qiagen). 5 ng of cDNA was added to each reaction, and qRT-PCR was performed using SYBR green assay (Qiagen). The data was then analyzed using the delta delta CT method and samples were paired within a culture. Genes were considered undetectable if the CT value was above 35.

**Conditioned Media**

Conditioned media was produced by culturing primary keratinocytes in KGM-2 media with or without EPX and its substrates $H_2O_2$ (100 μM) and $Br^-$ (100 μM), for 8 hours. Media was collected and centrifuged at 300 x g for 10 minutes to remove cells.

**Figures**

**Supplementary Table S1. Primer sequences used for qRT-PCR experiments.**

**Supplementary Table S2. Changes in cytokine expression in keratinocytes treated with EPX.** Keratinocytes were treated EPX and its substrates for 6 hours and RNA was harvested and analyzed using the Qiagen common cytokines RT2 profiler array. Gene expression is calculated as a fold change over its paired control sample. Each value is the average of 3 independent experiments, N/D = not detected.

**Supplementary Figure S1. Major basic protein is not required for inflammation and itch following exposure to TMA**. A) TMA induced itching was measured using time lapsed videography after the final TMA exposure. Quantification of itching of TMA treated ears in WT and MBP-1$^{-/-}$ mice were compared to the vehicle treated contralateral control ear from the same mouse. Data is expressed as a fold increase in scratching of TMA treated ear over contralateral control ear from same mouse. B) Sensitized WT and MBP-1 $^{-/-}$ mice were challenged on the ear with TMA, and ear thickness was measured on day 6 prior to the first challenge and on day 15. Change in ear thickness is expressed as difference in mm between two readings. E) Eosinophil were labeled in skin by staining eosinophils pink with a rat anti-eosinophil associated ribonuclease antibody.

**Supplementary Figure S2. Resorcinol dose dependently inhibits EPX activity.** Human EPX was incubated with its substrates in the presence of o-phenylenediamine (Sigma) and with increasing concentrations of resorcinol at room temperature for 15 minutes. The reaction was then stopped with 2N sulfuric acid and absorbance was read at 490 nm. EPX activity was compared with a negative control with no resorcinol.

**Supplementary Figure S3. Eosinophil peroxidase increases cytokine expression through a soluble mediator.** Cytokine gene expression in keratinocytes treated with conditioned media or EPX and its substrates alone for 4 hours. Conditioned media was generated by treating keratinocytes with EPX and its substrates for 8 hours, collecting the media, and centrifuging to remove cells. Gene expression is normalized to control treated cells. In the case of cells treated with conditioned media, the control is conditioned media from untreated cells.

Supplementary Table 1. Primer Sequences

| Gene | 5' Primer | 3' Primer |
|---|---|---|
| 18S | GTAACCCGTTGAACCCCATT | CCATCCAATCGGTAGTAGCG |
| TSLP | CTTGTCTCCTGAAAATCGAG | ATTCTGGAGATTGCATGAAG |
| TNF | CTGAACTTCGGGGTGATCGG | GGCTTGTCACTCGAATTTTGAGA |
| IL-1α | CGAAGACTACAGTTCTGCCATT | GACGTTTCAGAGGTTCTCAGAG |
| CSF2 | TCGTCTCTAACGAGTTCTCCTT | CGTAGACCCTGCTCGAATATCT |
| CSF3 | ATGGCTCAACTTTCTGCCCAG | CTGACAGTGACCAGGG3GAAC |
| sPLA2G1B | GTGTGGCAGTTCCGCAATATG | CCTGTCTAAGTCGTCCACTGG |
| sPLA2G2A | TGGCTCAATACAGGACCAAGG | GTGGCATCCATAGAAGGCATAG |
| sPLA2G2C | GCTGCCAACCCATCTTGAATG | CACAGACTGTTTGTCACACTCA |
| sPLA2G2D | TGCTGGCCGGTATAACTGC | CTGTGGCATCTTTGGGTTGC |
| sPLA2G2E | CCAGTGGACGAGACGGATTG | AGCAGCTCTCTTGTCACACTC |
| sPLA2G2F | GCCTCTCCCTCTAAAACCTCC | AGCACCAGTCTACCTCATCCA |
| sPLA2G3 | AGAGACCACAGGGCCATTAAG | GCTGTAGAATGACATGGTGCT |
| sPLA2G5 | CCAGGGGGCTTGCTAGAAC | AGCACCAATCAGTGCCATCC |
| sPLA2G7 | CTTTTCACTGGCAAGACACATC<br>T | CGACGGGGTACGATCCATTTC |
| sPLA2G10 | GTGCAGGTGTGACGAGGAG | CACTTGGGAGAGTCCTTCTCA |
| sPLA2G12<br>A | AGATAGACACGTACCTCAACGC | GCTGCACTTGTACTGGCAGA |
| LPAR1 | AGCCATGAACGAACAACAGTG | CATGATGAACACGCAAACAGTG |
| LPAR2 | ATGGTAGCTGTCTACACACGA | AACCGTCTTGACTAGGCTGAG |
| LPAR3 | CAAGCGCATGGACTTTTTCTAC | GAAATCCGCAGCAGCTAAGTT |
| LPAR4 | AGTGCCTCCCTGTTTGTCTTC | GCCAGTGGCGATTAAAGTTGTA<br>A |
| LPAR5 | ACCTGGACATGATGTTTGCCA | GAGACCAGTCGCCAATACCA |
| LPAR6 | ACGGGTGCATGTTCAGCAT | TGCCAGGTTAATCATGTACGTTG |

Supplementary Table 2.  Changes in cytokine expression in keratinocytes treated with EPX.

| Gene Name | Gene Symbol | Fold increase with EPX |
|---|---|---|
| Adiponectin C1Q and collagen domain containing | Adipoq | N/D |
| Aminoacyl tRNA synthetase complex-interacting multifunctional protein 1 | Aimp1 | 0.96 |
| Bone morphogenetic protein 1 | Bmp1 | 0.91 |
| Bone morphogenetic protein 2 | Bmp2 | 0.83 |
| Bone morphogenetic protein 3 | Bmp3 | 1.29 |
| Bone morphogenetic protein 4 | Bmp4 | 1.02 |
| Bone morphogenetic protein 5 | Bmp5 | N/D |
| Bone morphogenetic protein 6 | Bmp6 | 1.12 |
| Bone morphogenetic protein 7 | Bmp7 | 0.71 |
| CD40 ligand | Cd40lg | N/D |
| CD70 antigen | Cd70 | N/D |
| Ciliary neurotrophic factor | Cntf | 1.04 |
| Colony stimulating factor 1 (macrophage) | Csf1 | 1.75 |
| Colony stimulating factor 2 (granulocyte-macrophage) | Csf2 | 2.82 |
| Colony stimulating factor 3 (granulocyte) | Csf3 | 5.79 |
| Cardiotrophin 1 | Ctf1 | 1.01 |
| Fas ligand (TNF superfamily member 6) | Fasl | N/D |
| Fibroblast growth factor 10 | Fgf10 | N/D |
| Growth differentiation factor 15 | Gdf15 | 1.08 |
| Growth differentiation factor 2 | Gdf2 | N/D |
| Growth differentiation factor 5 | Gdf5 | N/D |
| Growth differentiation factor 9 | Gdf9 | 1.09 |
| Interferon alpha 2 | Ifna2 | N/D |
| Interferon alpha 4 | Ifna4 | N/D |
| Interferon beta 1 | Ifnb1 | N/D |
| Interferon gamma | Ifng | N/D |
| Interleukin 10 | Il10 | N/D |
| Interleukin 11 | Il11 | 0.70 |

| | | |
|---|---|---|
| Interleukin 12B | Il12b | N/D |
| Interleukin 13 | Il13 | N/D |
| Interleukin 15 | Il15 | 0.82 |
| Interleukin 16 | Il16 | 0.69 |
| Interleukin 17A | Il17a | N/D |
| Interleukin 17B | Il17b | 0.65 |
| Interleukin 17C | Il17c | 1.47 |
| Interleukin 17F | Il17f | N/D |
| Interleukin 18 | Il18 | 1.02 |
| Interleukin 19 | Il19 | 3.81 |
| Interleukin 1 alpha | Il1a | 1.93 |
| Interleukin 1 beta | Il1b | N/D |
| Interleukin 1 receptor antagonist | Il1rn | 2.28 |
| Interleukin 2 | Il2 | N/D |
| Interleukin 20 | Il20 | N/D |
| Interleukin 21 | Il21 | N/D |
| Interleukin 23 alpha subunit p19 | Il23a | 2.27 |
| Interleukin 24 | Il24 | 1.22 |
| Interleukin 25 | Il25 | N/D |
| Interleukin 27 | Il27 | N/D |
| Interleukin 3 | Il3 | N/D |
| Interleukin 4 | Il4 | N/D |
| Interleukin 5 | Il5 | N/D |
| Interleukin 6 | Il6 | N/D |
| Interleukin 7 | Il7 | 2.27 |
| Interleukin 9 | Il9 | N/D |
| Inhibin alpha | Inha | 1.29 |
| Inhibin beta-A | Inhba | 2.01 |
| Left right determination factor 1 | Lefty1 | 1.06 |
| Leukemia inhibitory factor | Lif | 1.94 |
| Lymphotoxin A | Lta | N/D |
| Lymphotoxin B | Ltb | 0.89 |
| Macrophage migration inhibitory factor | Mif | 0.86 |
| Myostatin | Mstn | N/D |
| Nicotinamide phosphoribosyltransferase | Nampt | 1.05 |
| Oncostatin M | Osm | N/D |
| Secretoglobin | Scgb3a1 | 0.96 |

| | | |
|---|---|---|
| Secreted phosphoprotein 1 | Spp1 | 1.29 |
| Transforming growth factor beta 1 | Tgfb1 | 1.45 |
| Transforming growth factor beta 2 | Tgfb2 | 1.56 |
| Thrombopoietin | Thpo | N/D |
| Tumor necrosis factor | Tnf | 2.54 |
| Tumor necrosis factor receptor superfamily member 11b (osteoprotegerin) | Tnfrsf11b | 1.32 |
| Tumor necrosis factor (ligand) superfamily member 10 | Tnfsf10 | N/D |
| Tumor necrosis factor (ligand) superfamily member 11 | Tnfsf11 | N/D |
| Tumor necrosis factor (ligand) superfamily member 12 | Tnfsf12 | N/D |
| Tumor necrosis factor (ligand) superfamily member 13 | Tnfsf13 | N/D |
| Tumor necrosis factor (ligand) superfamily member 13b | Tnfsf13b | N/D |
| Tumor necrosis factor (ligand) superfamily member 14 | Tnfsf14 | N/D |
| Tumor necrosis factor (ligand) superfamily member 15 | Tnfsf15 | 1.23 |
| Tumor necrosis factor (ligand) superfamily member 18 | Tnfsf18 | 1.56 |
| Tumor necrosis factor (ligand) superfamily member 4 | Tnfsf4 | 0.84 |
| Tumor necrosis factor (ligand) superfamily member 8 | Tnfsf8 | N/D |
| Tumor necrosis factor (ligand) superfamily member 9 | Tnfsf9 | 0.87 |
| Taxilin alpha | Txlna | 0.78 |
| Vascular endothelial growth factor A | Vegfa | 1.02 |

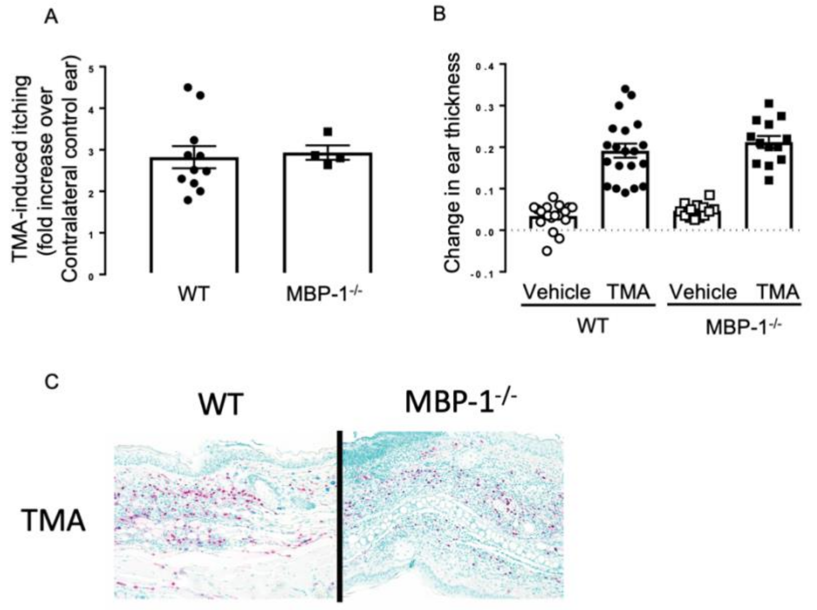


**Supplementary Figure S1**

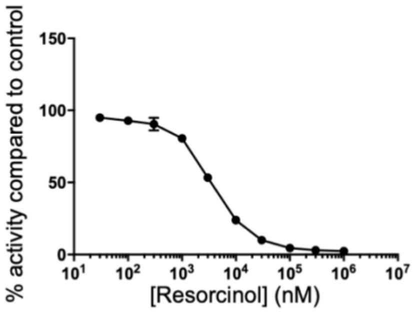


**Supplementary Figure S2**

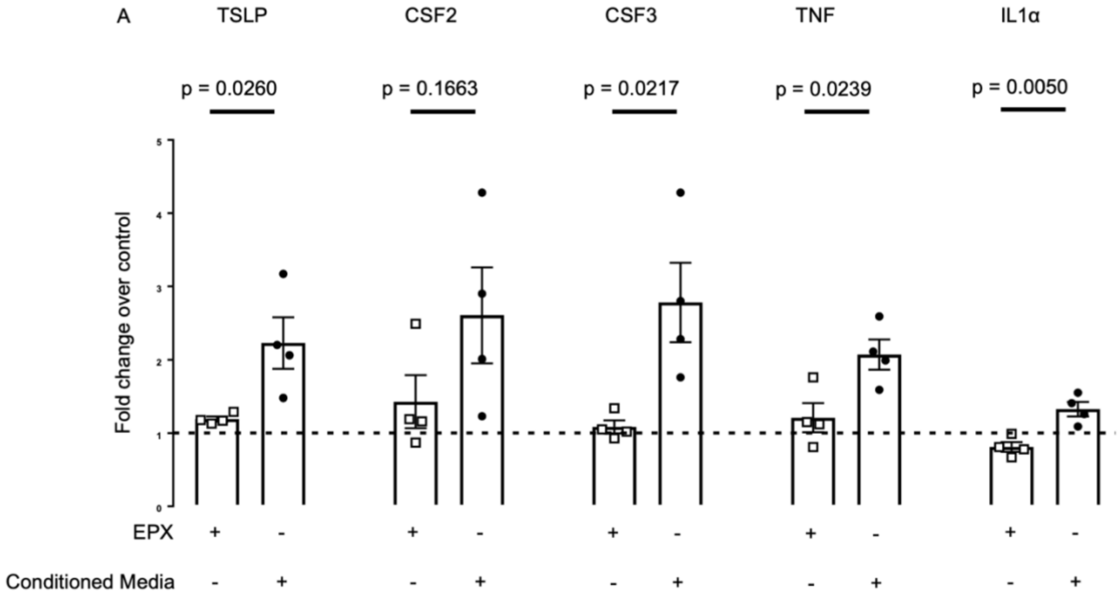


**Supplementary Figure S3**
